# The role of *fruitless* in specifying courtship behaviors differs across *Drosophila* species

**DOI:** 10.1101/2023.09.01.556001

**Authors:** Christa A. Baker, Xiao-Juan Guan, Minseung Choi, Mala Murthy

## Abstract

Sex-specific behaviors are critical for reproduction and species survival. The sex-specifically spliced transcription factor *fruitless* (*fru*) helps establish male courtship behaviors in invertebrates. Forcing male-specific *fru* (*fruM*) splicing in *Drosophila melanogaster* females produces male-typical behaviors, while disrupting female-specific behaviors. However, whether *fru*’s joint role in specifying male and inhibiting female behaviors is conserved across species is unknown. We used CRISPR/Cas9 to force FruM expression in female *D. virilis*, a species in which males and females produce sex-specific songs. In contrast to *D. melanogaster*, in which one *fruM* allele is sufficient to generate male behaviors in females, two alleles are needed in *D. virilis* females. *D. virilis* females expressing FruM maintain the ability to sing female-typical song as well as lay eggs, whereas *D. melanogaster* FruM females cannot lay eggs. These results reveal important differences in *fru* function between divergent species and underscore the importance of studying diverse behaviors and species for understanding the genetic basis of sex differences.

## Introduction

Sex-specific behaviors, such as reproduction, aggression, and parental care, are essential for species survival. Many of these innate behaviors are under genetic control in both vertebrates and invertebrates (*1*). In insects, sex-specific splicing of two transcription factors called *fruitless* (*fru*) and *doublesex* are responsible for establishing sexually differentiated neural circuitry and somatic tissue (*2–10*). In *Drosophila melanogaster*, in which *fru* function was first described (*11*), splicing in the male pattern results in a functional protein called FruM in a subset (∼2000, or ∼2%) of male neurons, whereas splicing in the female pattern results in transcripts that are degraded, leading to no functional protein (*3, 12, 13*). Sex-specific *fru* splicing has since been found across insect species (*14–18*) (but see (*19*)), but whether the role of *fru* in specifying sex-specific behaviors is conserved between *D. melanogaster* and other species remains an open question.

Knocking out FruM expression in male *D. melanogaster* eliminates their ability to engage in courtship behaviors directed toward a female (*3, 20*), and this function appears to be conserved in insects (*17, 21–24*) (but see (*25*)). Subsets of FruM-expressing neurons play distinct roles in producing male courtship behaviors in *D. melanogaster* (*5, 26–30*), and at least some of these neurons are conserved across *Drosophila* species despite divergence in courtship behaviors (*20, 31, 32*). A breakthrough in our understanding of *fru* function came from forcing male-specific *fru* splicing in female *D. melanogaster* (*3, 6*). This experiment was critical because *fruM* transcripts are also alternatively spliced at the 3’ end (Fig. 1A), giving rise to three isoforms differentially expressed across neurons: FruM-A, FruM-B, and FruM-C (*3, 30, 33, 34*). Females with male-specific splicing of *fruM* not only made FruM protein, but expressed the correct isoform in each cell type. Surprisingly, these females engaged in male courtship behaviors, with disruptions in their ability to produce female-specific behaviors - hence *fru* was considered a ‘switch gene’ for specifying sexually dimorphic behaviors (*3*). Here, we use a similar strategy (via CRISPR/Cas9 gene editing) to, for the first time, force FruM expression in females of another *Drosophila* species, and investigate whether this also results in male-typical courtship behaviors while impeding female-typical behaviors. In other words, does FruM expression in females result in different outcomes depending on the species context?

**Figure 1.**
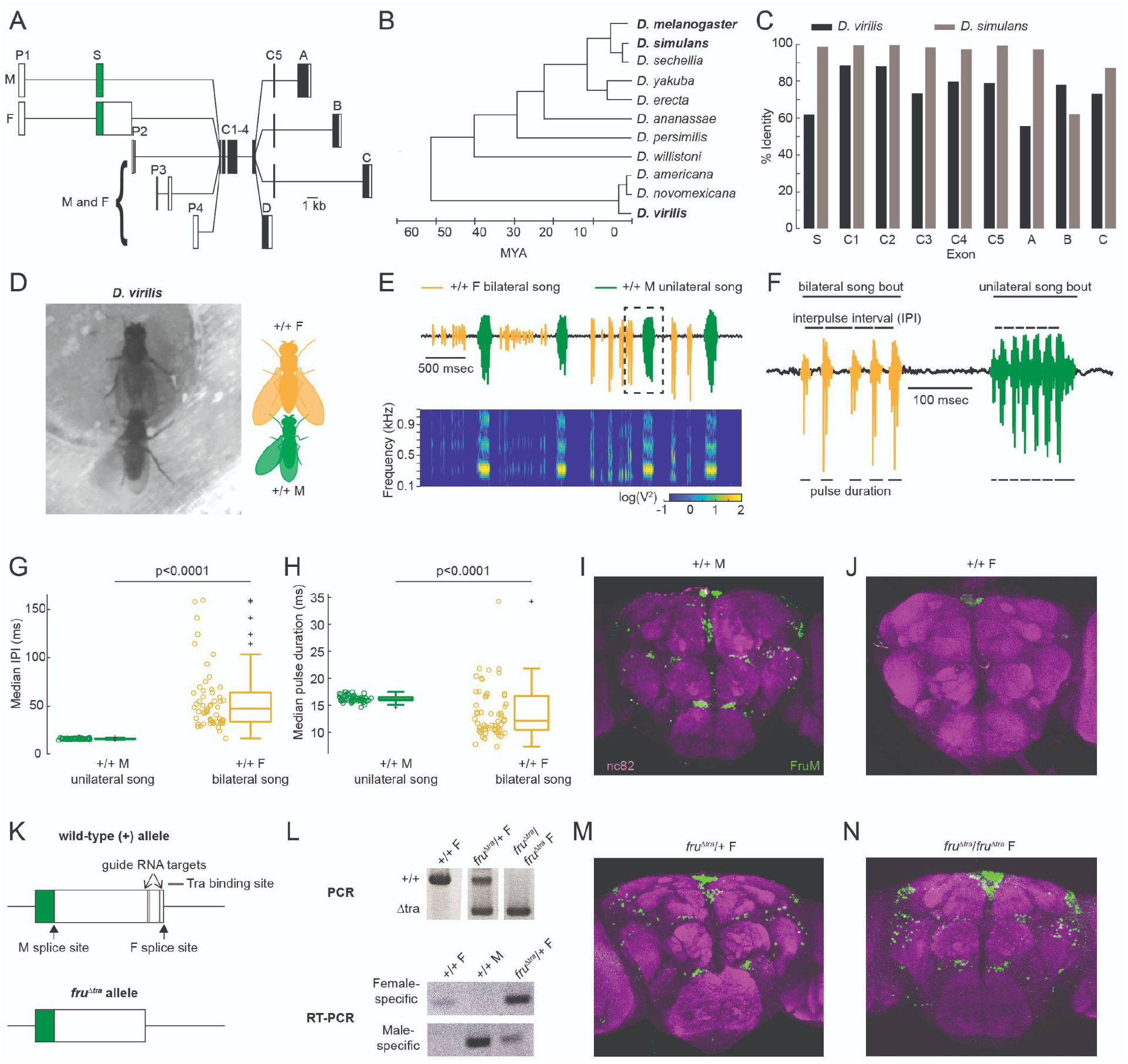
CRISPR-Cas9 gene editing of the *fruitless* (*fru*) gene results in expression of male-specific FruM in *D. virilis* female brains. (**A**) Transcripts resulting from alternative splicing of the *fru* gene. Filled and open boxes indicate coding and non-coding regions, respectively. P1-4 indicate promoters, S indicates the sex-specifically spliced exon, C1-5 indicate exons common to most *fru* transcripts, and A-D indicate alternative 3’ exons. Adapted from (*3*). (**B**) *Drosophila* phylogeny. Adapted from (*37*). (**C**) Comparison of the *fru* exon (coding regions only) nucleotide sequences. Percent identity is reported relative to *D. melanogaster*. (**D**) Video still (left) and schematic (right) of *D. virilis* courtship duets. Wild-type (+/+) *D. virilis* males and females both sing during courtship, with females using bilateral wing vibration and males using unilateral wing vibration. (**E**) Four-second microphone recording (top) and spectrogram (bottom) of duet example. Unilateral song is stereotyped in temporal pattern and frequency, whereas bilateral song is more variable. Songs were automatically segmented using a convolutional neural network (Fig S3; see Methods). (**F**) Close-up of the microphone recording outlined in (**E**). We measured the interpulse intervals (IPIs) and pulse durations as indicated. (**G-H**) IPI (**G**) and pulse durations (**H**) of unilateral and bilateral song. Each dot represents the median IPI from one fly. Statistical tests were Wilcoxon rank sum tests with Bonferroni correction. n=55 flies in both groups. (**I-J**) Antibody staining for FruM (green) and bruchpilot (nc82; magenta) in *D. virilis* +/+ male (**I**) and +/+ female (**J**) brains. The ocelli in both sexes are immunopositive for FruM. (**K**) Schematic of the S-exon (top) and the result of CRISPR-Cas9-mediated removal of the Transformer (Tra) binding sites (*fru*^*∆ tra*^) (bottom). (**L**) (top) PCR genotyping of *fru*^*∆ tra*^ mutants using primers flanking the CRISPR guide RNA targets. Heterozygotes have both a +/+ and a shorter mutant allele (middle), whereas homozygous mutants have only the mutant allele (right). (bottom) RT-PCR using *D. virilis* female- and male-specific primers confirm that the brains of *fru*^*∆ tra*^*/+* females contain male *fru* transcripts. (**M-N**) FruM antibody staining in *D. virilis fru*^*∆ tra*^*/+* (**M**) and *fru*^*∆ tra*^*/fru*^*∆ tra*^ (**N**) female brains.

We selected *D. virilis*, which has male-specific FruM expression (*35, 36*) but which diverged from *D. melanogaster* almost 60 million years ago (Fig. 1B) (*37*), and shows only ∼50-90% sequence identity (Fig. 1C). Importantly for our experiments, *D. virilis* produces markedly divergent courtship behaviors compared to *D. melanogaster*. Courtship in *D. melanogaster* consists of the male pursuing the female, while tapping, licking, and singing to her with unilateral wing extensions (*38*). Females in turn arbitrate mating decisions by slowing down and allowing copulation when receptive, and then laying eggs. In the vast majority of drosophilid species with courtship songs, it is the male who sings to the female (*39*). Females have been reported to sing back to males in just a few species, all within the *D. virilis* group (*40*). *D. virilis* males use unilateral wing vibration while females use bilateral wing vibration to produce sex-specific pulse songs during acoustic duets (Fig 1D-E) (*41*). Males can also sing a female-like bilateral song on the infrequent occasions when they are courted by another male (*41*), demonstrating that song choice is context-dependent in males. In contrast, females do not naturally produce the male-typical unilateral song. Therefore, unilateral song is male-specific, whereas bilateral song appears to be sexually monomorphic in *D. virilis*.

Here we present evidence suggesting evolutionary divergence in the role of *fru* between *D. virilis* and *D. melanogaster*. Although FruM expression in *D. virilis* females results in male courtship behaviors, including unilateral song production, these effects require two alleles of *fruM* in *D. virilis* but just one allele in *D. melanogaster* (*3*). *D. virilis* females expressing FruM maintain the ability to sing female-like bilateral song, although bilateral song features are changed - such a function was not possible to query in *D. melanogaster*, since those females do not sing. Similar to *D. melanogaster*, FruM expression reduced receptivity in *D. virilis* females, but in contrast to *D. melanogaster*, FruM expression did not eliminate egg-laying in *D. virilis* females. *D. virilis*, but not *D. melanogaster*, FruM females also exhibited male-directed aggression. These results highlight the value of comparing diverse behaviors and species in evaluating the role of important ‘switch genes’ involved in behavioral specification.

## Results

In *D. virilis* duets, two key features distinguish male-typical unilateral from female-typical bilateral song: the time between successive pulses, called interpulse intervals (IPIs), and the duration of each pulse (Fig 1F). These features are stereotyped within and across males, but are more variable in females (Fig 1G-H). In addition to unilateral song, *D. virilis* males can sing the female-typical bilateral song if they are courted by another male (Fig S1A-C) (*41*), though male-directed courtship occurs less frequently than female-directed (Fig S1D). Male and female bilateral song is similar (Fig S1E-G), suggesting that *D. virilis* may have two song circuits: one sexually monomorphic circuit for bilateral song and one dimorphic circuit for unilateral song. The role of *fru* in establishing either of these song circuits is unknown.

### Transformer binding site removal results in FruM expression in *D. virilis* female brains

Similar to *D. melanogaster*, FruM expression in *D. virilis* is male-specific (Fig. 1I-J) (*35, 36*). To understand the role FruM plays in specifying *D. virilis* courtship behaviors, we forced *fruM* splicing in female brains by removing the Transformer (Tra) binding sites from the sex-specifically spliced S exon (Fig 1K) using CRISPR-Cas9, which in *D. melanogaster* results in splicing at the male site (*3*). This mutation is equivalent to the *fru*^*∆ tra*^ mutation in *D. melanogaster* (*3*). PCR (Fig 1L, top) and sequencing confirmed that our mutagenesis resulted in removal of a portion of the S-exon containing the Tra binding sites. RT-PCR against female- and male-specific *fru* transcripts confirmed that *fru*^*∆ tra*^/+ female brains contained both versions (Fig 1L, bottom), demonstrating that Tra binding site removal was sufficient to produce male-specific *fru* transcripts. We then validated these results with antibody staining for FruM (see Materials and Methods), and found that females carrying the *fru*^*∆ tra*^ mutation had FruM expression in subsets of neurons (Fig 1M-N) that overall was consistent with the +/+ male expression pattern (Fig 1I).

### *ruM* is haploinsufficient for male courtship behaviors in *D. virilis* females

The effects of FruM expression in *D. melanogaster* females requires only one copy of the *fru*^*∆ tra*^ allele (*3*), a pattern known as haplosufficiency or dominance. This suggests that the amount of FruM transcription factor resulting from one *fru*^*∆ tra*^ allele is sufficient to produce masculinized neural circuitry in females. To determine whether *fruM* is also dominant in *D. virilis*, we paired a single *fru*^*∆ tra*^/+ female with a wild-type female (Fig 2A) and quantified courtship behaviors. *D. virilis* courtship consists of the male orienting to the female, rubbing the underside of her abdomen with his front tarsi, licking her genitalia, and singing with unilateral wing extensions (*41*). We found that *D. virilis fru*^*∆ tra*^/+ females exhibited very little male-specific courtship behavior (light blue in Fig 2B-D), and no unilateral song in pairings with wild-type females (Fig 2D). These results differ from those in *D. melanogaster* females, in which one allele of *fru*^*∆ tra*^ leads to male courtship behaviors (*3*).

**Figure 2.**
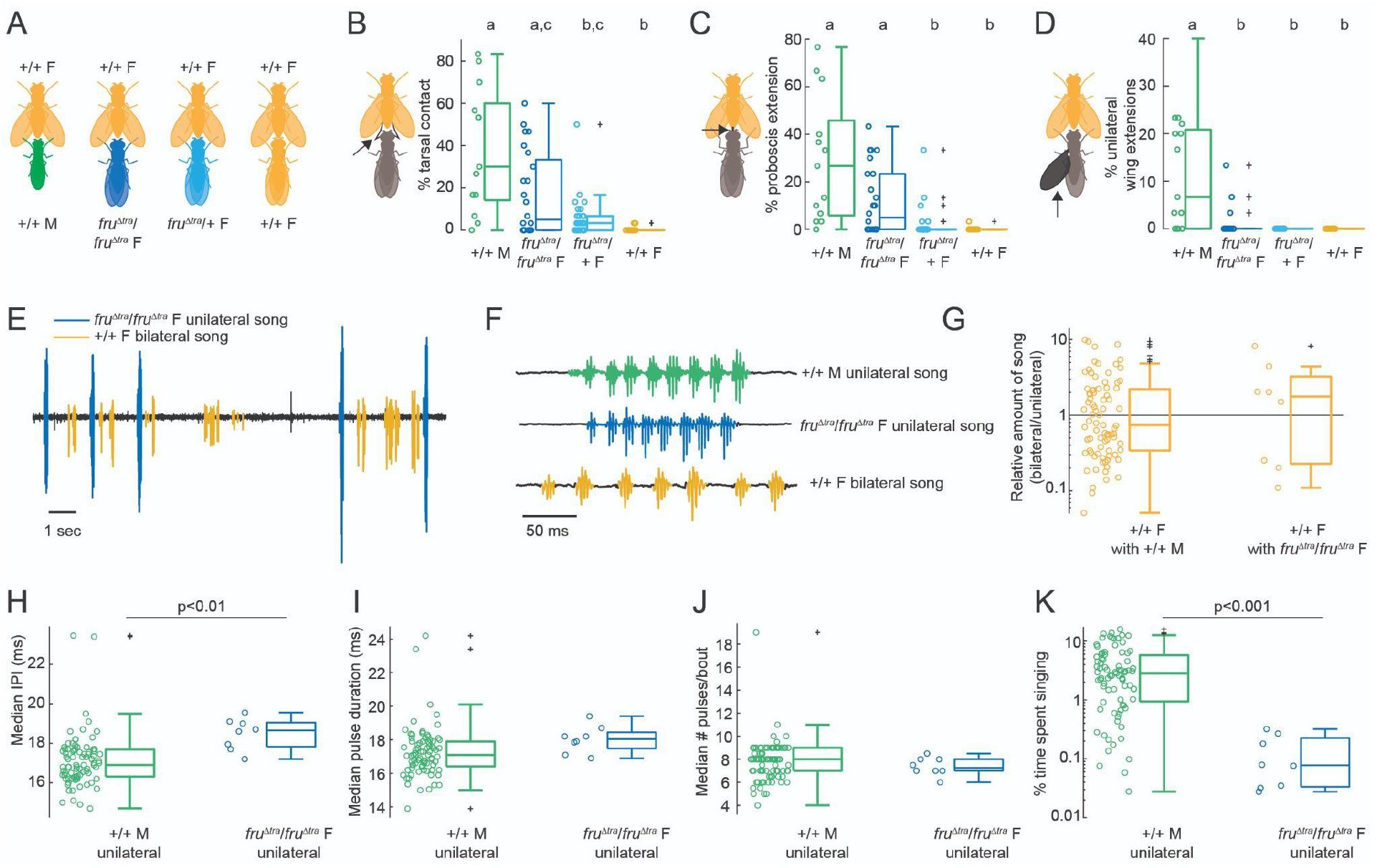
*D. virilis fru*^*∆ tra*^/*fru*^*∆ tra*^ females are capable of producing male-typical courtship behaviors directed toward +/+ females. (**A**) To test whether FruM specifies male courtship behaviors in *D. virilis*, we paired single *fru*^*∆ tra*^/*fru*^*∆ tra*^ and *fru*^*∆ tra*^/*+* females with a +/+ female. +/+ males and +/+ females (siblings to the *fru*^*∆ tra*^ females) each paired with a +/+ female served as controls. (**B-D**) % of manually scored bins containing instances where the experimental fly contacted the +/+ female with their front tarsi (arrow in pictogram), extended their proboscis (**C**, arrow), and extended wings unilaterally (**D**, arrow). Each dot represents the value for a single fly. Statistical significance was determined by Kruskal-Wallis tests followed by pairwise Wilcoxon rank sum tests with Bonferroni correction. n=14, 24, 39, 14 flies. (**E**) 15 sec audio recording of a duet between a *fru*^*∆ tra*^/*fru*^*∆ tra*^ female and a +/+ female. (**F**) Close-up of the waveforms of +/+ male unilateral, *fru*^*∆ tra*^/*fru*^*∆ tra*^ female unilateral, and +/+ female bilateral songs. (**G**) Amount of bilateral song produced by the +/+ female normalized by the amount of unilateral song produced by the +/+ male (left) and *fru*^*∆ tra*^/*fru*^*∆ tra*^ female (right). Each dot represents one pair of flies. (**H-K**) Median IPI (**H**), median pulse duration (**I**), median number of pulses per bout (**J**), and percent time each fly spent singing unilateral song (**K**). n=83 and 8 flies in (**G-K**). Statistical tests were Wilcoxon rank sum tests with Bonferroni correction.

In contrast to *D. melanogaster* (*3*), *D. virilis fru*^*∆ tra*^/+ females produced offspring after copulating with males; we found that *fru*^*∆ tra*^/*fru*^*∆ tra*^ females courted wild-type females (Movie S1), with no significant differences in the amount of tarsal contact or proboscis extension compared to wild-type males (Fig 2B-C). We also observed unilateral wing extension in a few *fru*^*∆ tra*^/*fru*^*∆ tra*^ females (Fig 2D), suggesting that *fru*^*∆ tra*^/*fru*^*∆ tra*^ females may indeed sing unilateral song when paired with a female.

To quantify song production, we built a new *D. virilis* song segmenter consisting of two convolutional neural networks (see Materials and Methods): one trained to distinguish among four signal classes (unilateral song, bilateral song, overlap, no song) (Fig S2A), and a second trained to distinguish between two signal classes (bilateral song and no song) (Fig S2B).

Combining the output of the two networks (Fig S2C) resulted in high precision and sensitivity for detecting both unilateral and bilateral song, with equally good performance across genotypes (Fig S2D-F). The performance of this segmenter is superior to that previously developed for *D. virilis* duets (*41*).

We found unilateral song in 8/24 (33%) pairings between *fru*^*∆ tra*^/*fru*^*∆ tra*^ *D. virilis* females and wild-type females. This song alternated with bilateral song from the wild-type female (Fig 2E), and occurred in stereotyped bouts that looked similar to wild-type male unilateral song (Fig 2F). We visually confirmed that the unilateral song occurred when the *fru*^*∆ tra*^/*fru*^*∆ tra*^ female was indeed performing unilateral wing extensions, and that the bilateral song was produced solely by the wild-type female (Movie S1). Wild-type females sang just as much bilateral song with *fru*^*∆ tra*^/*fru*^*∆ tra*^ females as with wild-type males (Fig 2G). *fru*^*∆ tra*^/*fru*^*∆ tra*^ unilateral song had short IPIs in line with those of wild-type males, although the IPI was modestly but significantly increased by 1-2 ms (Fig 2H). There was no difference in pulse duration (Fig 2I) or the number of pulses per bout (Fig 2J) between the unilateral song from *fru*^*∆ tra*^/*fru*^*∆ tra*^ females and wild-type males.

However, we found that *fru*^*∆ tra*^/*fru*^*∆ tra*^ females produced almost an order of magnitude less unilateral song than wild-type males (Fig 2K). Together, these results reveal that, while FruM specifies male-typical unilateral song in *D. virilis*, it is not sufficient (even with two alleles) to produce male-typical levels of courtship drive. Since two alleles of *fruM* are required for female *D. virilis* to produce male-like courtship behaviors, *fruM* is haploinsufficient for male-typical behaviors, including unilateral song, in female *D. virilis* (in contrast with *D. melanogaster*). Since FruM is a transcription factor involved in the development of sexually dimorphic neural circuitry, this finding suggests that although *fru*^*∆ tra*^/*+* and *fru*^*∆ tra*^/*fru*^*∆ tra*^ *D. virilis* females developed FruM+ neurons (Fig. 1M-N), there may be important differences in FruM+ neuron number, morphology, function, and/or connectivity patterns dependent on allele number.

We next returned to *D. melanogaster* to re-investigate *fruM* allele number and song production. We conducted single-pair courtship experiments using two *D. melanogaster fru*^*∆ tra*^ genotypes (Fig S3A): *fru*^*∆ tra*^/+ to match the genotype of the *D. virilis fru*^*∆ tra*^*/+* females, and *fru*^*∆ tra*^/*fru*^*4-40*^ (where *fru*^*4-40*^ is a fruM-null mutation) (*12*), to compare with previous experiments (*3, 42*). Both *fru*^*∆ tra*^/+ (Movie S2) and *fru*^*∆ tra*^/*fru*^*4-40*^ *D. melanogaster* females courted wild-type females. Whereas both *fru*^*∆ tra*^/+ and *fru*^*∆ tra*^/*fru*^*4-40*^ females engaged in tapping and unilateral wing extensions directed toward a wild-type female (Fig S3B-D), the amount of these behaviors were significantly less than those produced by wild-type males (Fig S3C-D). Consistent with prior work (*42*), we found that while *fru*^*∆ tra*^ females extended their wings, this did not produce bonafide song (Fig S3E-F).

Taken together, our results reveal two key species differences in the role of FruM in specifying male courtship behaviors: 1) FruM is haploinsufficient to produce male courtship in *D. virilis* females but not in *D. melanogaster* females; 2) two alleles of *fruM* produce male-typical unilateral song in female *D. virilis*, whereas the effects of two alleles of *fruM* cannot be tested in female *D. melanogaster* as one allele renders females infertile (*3*).

### Removing a *fruM* allele in *D. virilis* males has no effect on courtship behaviors

The haploinsufficiency of *fruM* to produce unilateral song in *D. virilis* females raised the question of whether *fruM* is also haploinsufficient in males. Wild type males make (via splicing) two functional copies of FruM RNA in each cell, one from each allele. What happens to male behavior if we remove one of these copies? We generated males lacking one copy of *fruM* by removing the S-exon (Fig S4A) via CRISPR-Cas9 and confirming the removal with PCR (Fig S4B-C) and sequencing. We refer to these males as -/+. We then paired -/+ males with wild-type females (Fig S4D) and found that -/+ males robustly courted females. There was a modest reduction in tarsal contact by -/+ males compared to wild-type males (Fig S4E), but no differences in the amount of proboscis extension (Fig S4F) or unilateral wing extension (Fig S4G). Wild type females duetted with -/+ males, and the waveforms of -/+ male unilateral song appeared similar to wild-type male unilateral song (Fig S4H-J). We found no differences in the quantitative parameters of -/+ unilateral song or in the amount or timing of unilateral song (Fig S4K-O). -/+ males were also as likely to copulate as wild-type males (Fig S4P). Taken together, these results demonstrate that *fruM* is haploinsufficient for the production of male courtship behaviors, including unilateral song, *only* in the female context in *D. virilis*.

### FruM disrupts female receptivity but not egg-laying in *D. virilis*

FruM expression in female *D. melanogaster* not only results in male-typical courtship behaviors, but also interferes with female-typical courtship behaviors, such as egg-laying and receptivity (*3*). In contrast to *D. melanogaster, D. virilis fru*^*∆ tra*^*/+* females maintained their ability to lay eggs, as almost 50% of *fru*^*∆ tra*^*/+* females that copulated produced offspring (Fig. S5A). In our single-pair courtship assays, the copulation rate of *D. virilis fru*^*∆ tra*^*/+* females was about 40% that of wild-type females (Fig. S5B). In a striking similarity between species, the copulation rate of *D. melanogaster fru*^*∆ tra*^ females was also 35-40% that of wild-type females (Fig. S5C). These results point to divergent effects of FruM expression on egg-laying but conserved effects on female receptivity between the two species.

Adding a second allele of *fru*^*∆ tra*^ to *D. virilis* females completely eliminated copulation (Fig. S5B). This was not due to reduced attractiveness to males, as males directed equal amounts of courtship behaviors toward females of all genotypes (Fig S5D-F). Therefore, FruM effects on female receptivity in *D. virilis* depend on the number of *fruM* alleles.

### FruM alters female-typical bilateral song features and amount

The effects of FruM expression on female singing behavior has not previously been tested in any species. Since *D. virilis* bilateral song is sexually monomorphic (Fig S1) (*41*), we expected FruM expression in females to have no effect on bilateral song production. In pairings of *fru*^*∆ tra*^/+ and *fru*^*∆ tra*^/*fru*^*∆ tra*^ *D. virilis* females with wild-type males (Fig 3A), *fru*^*∆ tra*^ females readily duetted with wild-type males (Fig 3B; Movie S3). The waveform of *fru*^*∆ tra*^ bilateral song looked similar to wild-type female bilateral song (Fig 3C). However, one allele of *fru*^*∆ tra*^ resulted in increased IPIs by about 10 msec relative to wild-type females (Fig 3D). FruM expression also led to longer pulses (Fig 3E), and shorter response times (Fig 3F), regardless of the number of *fru*^*∆ tra*^ alleles.

**Figure 3.**
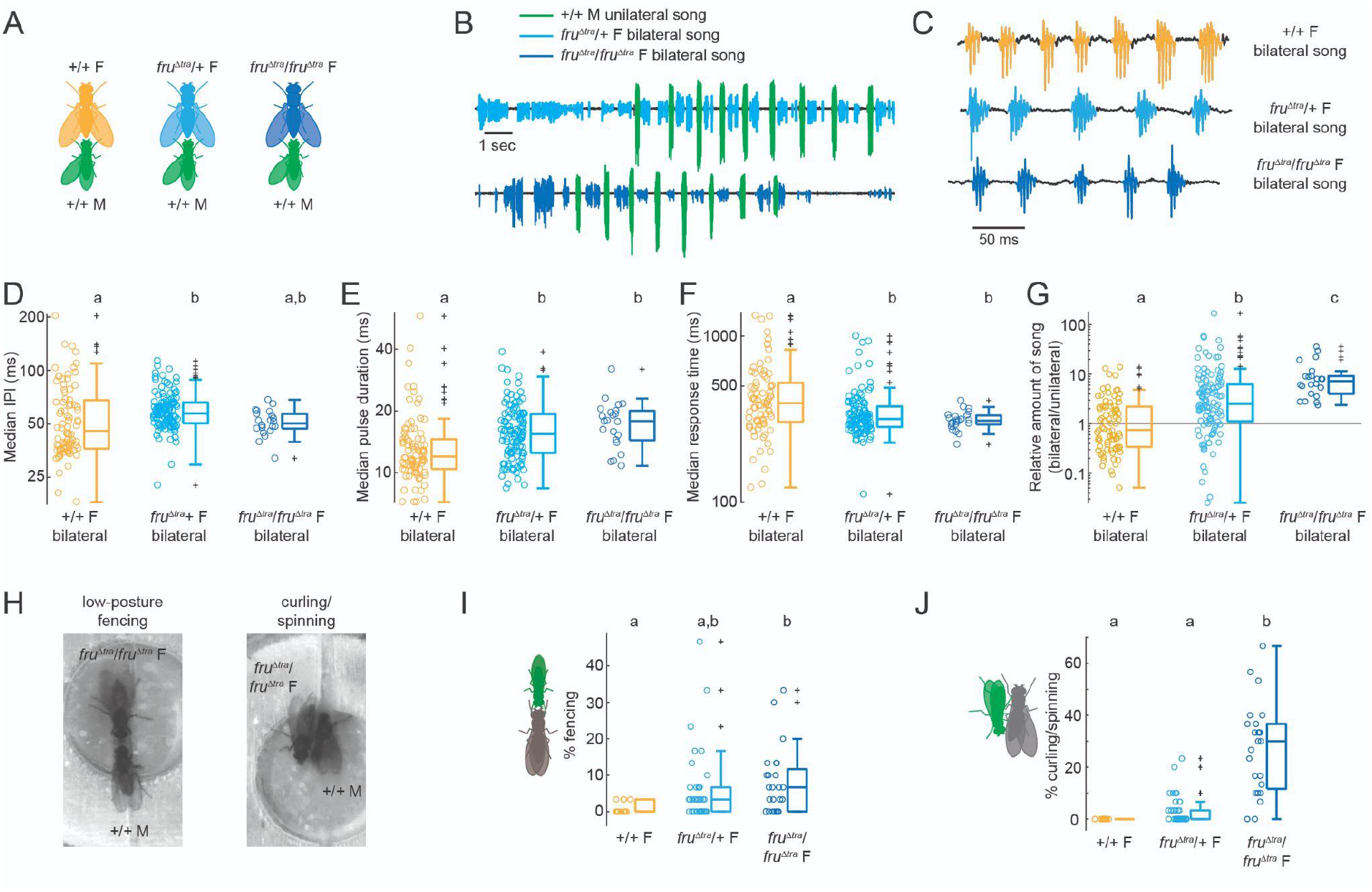
FruM expression in *D. virilis* females has effects on bilateral song and aggression. (**A**) To determine whether FruM expression affected bilateral song production, we paired single *fru*^*∆ tra*^*/+* and *fru*^*∆ tra*^*/fru*^*∆ tra*^ *D. virilis* females with a +/+ male. Single +/+ females (siblings of the *fru*^*∆ tra*^ females) paired with a +/+ male served as controls. (**B**) 15 sec microphone recordings of duets between *fru*^*∆ tra*^ females singing bilateral song and +/+ males singing unilateral song. (**C**) Close-up of bilateral song waveforms. (**D-G**) Bilateral song median IPI (**D**), pulse duration (**E**), response time (delay between onset of unilateral bout and center of first following bilateral pulse) (**F**), and amount relative to unilateral song (**G**). Each dot represents one fly. n=83, 114, 22 flies. Statistical tests were Kruskal-Wallis tests followed by pairwise Wilcoxon rank sum tests with Bonferroni correction. (**H**) Video stills of low-posture fencing (left) and curling/spinning (right).(**I-J**) Percentage of bins with fencing (**I**) and curling/spinning (**J**). Each dot represents one fly. n=13, 38, 24 flies.

Surprisingly, we also found that FruM expression increased the amount of bilateral song, with *fru*^*∆ tra*^/*fru*^*∆ tra*^ females singing almost an order of magnitude more bilateral song than wild-type females when paired with a wild-type male (Fig 3G). Increased bilateral song in *fru*^*∆ tra*^ females is consistent with the increased bilateral song produced by males relative to wild-type females when courted by another male (Fig S1H), suggesting that an upregulation of bilateral song may be a consequence of FruM-induced masculinization. Taken together, our results show that although bilateral song is sexually monomorphic, at least some of the underlying neural circuitry may be regulated by *fru*.

### Homozygous *fru*^*∆ tra*^ *D. virilis* females display male-directed aggression

We observed two types of male-directed aggressive behaviors from *D. virilis fru*^*∆ tra*^ females (Fig 3H) - aggressive behaviors were not reported in *fru*^*∆ tra*^ *D. melanogaster* females (*3, 6*). In one behavior, the *fru*^*∆ tra*^ female and male face each other and extend their front tarsi toward one another (Movie S4), similar to previously described”low-posture fencing” (*43*). In the second behavior, the *fru*^*∆ tra*^ female curls her abdomen toward the male’s head, similar to previously described curling (*44*). While the female is curling, the male seems to still try to tap and lick her abdomen, which results in the two flies spinning together in a circle (Movie S5). These aggressive behaviors were interspersed with duetting - duetting would often immediately precede and/or follow an aggressive bout. The amount of fencing and curling/spinning behaviors were dependent on the number of *fru*^*∆ tra*^ alleles in females (Fig 3I-J).

*fru*^*∆ tra*^*/fru*^*∆ tra*^ *D. virilis* females were the most likely to produce aggression (Fig 3I-J), and were also least likely to copulate (Fig S5B). However, we found no difference in the amount of aggressive behaviors in copulated vs. non-copulated *fru*^*∆ tra*^/+ females (Fig S5G-H). These behaviors do not seem to be typical reactions to courtship of an unreceptive female, since non-copulating wild-type females did not engage in these behaviors (Fig 3I-J). We also did not observe aggressive behaviors in pairings between two wild-type males (dark green in Fig S4Q-R).

We observed spinning between a handful of -/+ males and wild-type males, but not fencing (light green in Fig S4Q-R), suggesting that these behaviors are dependent on the number of *fruM* alleles in *D. virilis* females but not males. Therefore, FruM appears to have divergent effects on aggression in *D. virilis* vs. *D. melanogaster* females.

### *fruM* effects on courtship and aggression depend on allele number in female, but not male, *D. virilis*

So far, our results suggest that, in female *D. virilis*, FruM effects on both unilateral and bilateral song production as well as aggression depend on the number of *fruM* alleles. To clarify the effects of FruM in both sexes, we performed principal components analysis (PCA) on courtship and aggressive behaviors exhibited by the same fly paired with a wild-type female vs. a wild-type male (Fig 4A). These results show that, compared to wild-type females, which do not produce any aggressive or male-typical courtship behaviors, adding one copy of *fru*^*∆ tra*^ to females caused movement primarily along PC2 (Fig 4A), which correlated positively with both aggression (curling/spinning) and courtship (tarsal contact) directed toward a wild-type male. A second *fru*^*∆ tra*^ allele caused females to also move along PC1 (Fig 4A), which correlated positively with courtship (tarsal contact and proboscis extension) directed toward a wild-type female. Only a few of the *fru*^*∆ tra*^/*fru*^*∆ tra*^ females overlapped in PCA space with wild-type males, suggesting that FruM, while sufficient to produce male courtship behaviors including unilateral song in some *D. virilis* females, is not sufficient to fully masculinize females. In contrast to the effects of *fruM* allele number in females, removing one *fruM* allele in males had no effect on courtship or aggression (Fig 4A). These results lend further support to the conclusion that *fruM* has allele-number-dependent effects in *D. virilis* females but not males.

**Figure 4.**
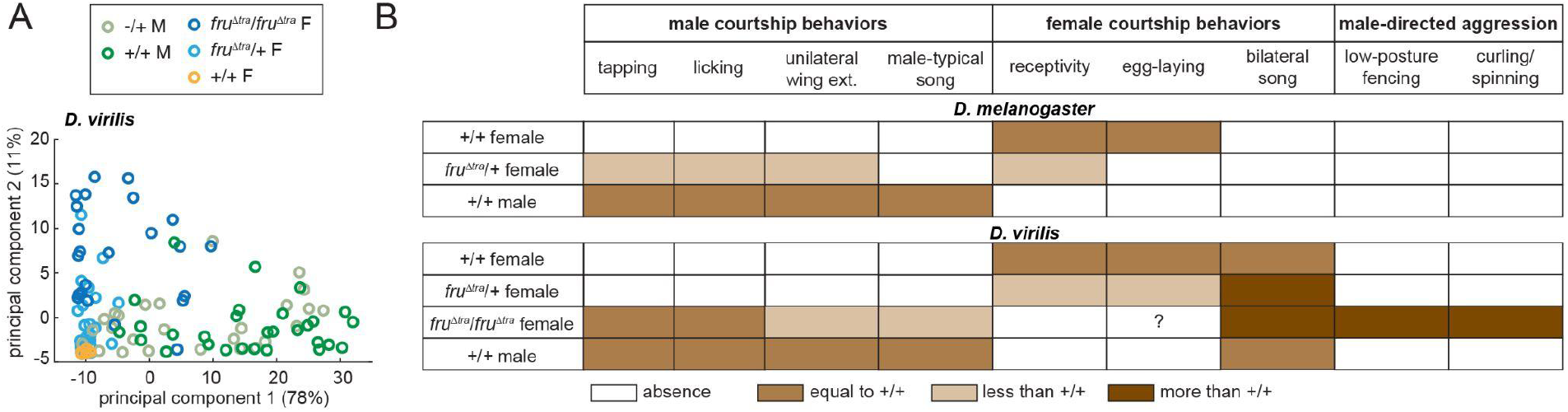
FruM effects on *D. virilis* courtship and aggressive behaviors depend on allele number in females. (**A**) Principal components analysis (PCA) on all scored courtship and aggressive behaviors exhibited by a single fly when paired with a +/+ female and (separately) a +/+ male. PC1 positively correlated with tarsal contact and proboscis extension directed toward a +/+ female, and PC2 positively correlated with curling/spinning and tarsal contact, both directed toward a +/+ male. Each dot represents one fly. n=26 -/+ M, 29 +/+ M, 24 *fru*^*∆ tra*^*/fru*^*∆ tra*^ F, 39 *fru*^*∆ tra*^*/+* F, 14 +/+ F flies. (**B**) Table summarizing the effects of FruM expression in *D. melanogaster* (top) and *D. virilis* (bottom). Colors indicate the amount of each behavior.

## Discussion

Here we provide evidence for divergence in the function of *fru* between two *Drosophila* species. Despite conservation of sex-specific splicing and expression of FruM in *D. virilis*, the function of FruM in specifying male, while disrupting female, courtship behaviors is different from *D. melanogaster*. We were able to uncover these differences by using CRISPR/Cas9 to force native male-specific *fruM* splicing in female *D. virilis*, which had previously been accomplished only in *D. melanogaster*. One of the most striking species differences we found is that *D. virilis*, but not *D. melanogaster*, females expressing FruM maintain their fecundity (Fig. 4B), enabling us to test the effects of one vs. two alleles of *fru*^*∆ tra*^ on behaviors in *D. virilis*, but not in *D. melanogaster*.

This revealed that FruM allele number in females (but not in males) affected the phenotypes observed. In contrast to *D. melanogaster*, in which one copy of *fru*^*∆ tra*^ is sufficient to produce some male courtship behaviors, two copies were needed in *D. virilis* (Fig. 4B). These male courtship behaviors were accompanied by male-typical song production in female *D. virilis* but not *D. melanogaster* (Fig. 4B) (*3, 42*). FruM expression in female *D. virilis* also had opposing effects on receptivity and bilateral (female-typical) song amount (Fig. 4B); effects on female song production could not be assayed in *D. melanogaster*, since those females do not sing. That FruM expression does not inhibit egg-laying or bilateral song production in *D. virilis* is different from the role of FruM in suppressing female behaviors in *D. melanogaster* and *Aedes aegypti* (*3, 21*). Finally, *D. virilis* females expressing FruM exhibited male-directed aggression, whereas *D. melanogaster* females did not (Fig. 4B).

### Differences between behaviors of *fru*^*∆ tra*^/*+* and *fru*^*∆ tra*^/*fru*^*∆ tra*^ *D. virilis* females

In *D. virilis*, we found differences in the amount of male courtship behaviors, bilateral song, aggression, and receptivity between females that were heterozygous vs. homozygous for *fru*^*∆ tra*^. This pattern is suggestive of haploinsufficiency, in which one copy of a gene is insufficient for a particular phenotype. Haploinsufficiency is not uncommon for transcription factors. For instance, FruM is haploinsufficient for pheromone responses in Or47b olfactory receptor neurons in male *D. melanogaste*r (*45*). In mice, haploinsufficiency of the transcription factor *Six3* disrupts male reproduction by impeding development of the main olfactory epithelium (*46*). In human sex determination, multiple transcription factors display haploinsufficiency leading to sex reversals (*47*). Causes of haploinsufficiency are hypothesized to include insufficient amounts of protein product arising from a single gene copy, stoichiometric disruptions of protein complexes, and more recently, a narrow range of tolerable protein expression levels (*47, 48*). Since FruM effects on cellular masculinization are cell-intrinsic (*49, 50*), a change in dosage-sensitivity and/or stoichiometry in just a small number of cells could account for FruM-dose-dependence in *D. virilis* compared to *D. melanogaster*. Therefore, *D. virilis* may represent an interesting case for investigating factors contributing to sex-specific transcription factor haploinsufficiency.

We found that *fruM* haploinsufficiency was specific to the female context in *D. virilis* - normal male courtship behavior was observed with just one allele of *fruM* (Fig. S4). Another transcription factor called *doublesex* (*dsx*) is also sex-specifically spliced in *D. melanogaster*, and, in contrast to *fru*, produces functional protein in males (DsxM) and females (DsxF) (*2*). Dsx is expressed in subsets of neurons that play key roles in both male- and female-specific behaviors (*5, 51–54*), and is co-expressed with FruM in some cell types in male brains (*5, 33, 42, 55*).

Females generally have fewer numbers of *dsx*+ and *fru*+ neurons than males due to DsxF-dependent cell death (*56*) and FruM-dependent inhibition of cell death (*57*). Therefore, while the presence of FruM in female *Drosophila* brains sufficiently masculinizes some cell types (*58*), DsxF may act to limit the extent of this masculinization. Our finding of fruM haploinsufficiency in *D. virilis* but not *D. melanogaster* females suggests that DsxF may exert different effects across species.

### Importance of analyzing the role of switch genes in other species

Findings in other insect species of sex-specific *fru* splicing and/or disruption of male courtship behaviors after *fruM* knockdown led to the hypothesis that *fru*’s role as a sex-determination switch gene was highly conserved (*14, 15, 17, 18, 20, 21, 23, 24, 59*). This hypothesis was also supported by findings that inserting *fru* genes from drosophilid species with divergent courtship behaviors into *D. melanogaster* males recapitulated *D. melanogaster* behaviors, instead of phenocopying each species’ own behaviors (*60*), which suggested that divergence in FruM downstream targets likely contributes to specifying species-specific behaviors. However, in most species previously studied, males and females engage in dramatically different behaviors during courtship. By choosing a species in which the sexes produce a similar behavior, ie, singing, we uncovered evolutionary divergence in *fru* function, despite conservation of sex-specific FruM expression (Fig 1I-J) (*35, 36*). Additional sex-determination switch genes in *D. melanogaster* include *dsx* and three genes upstream of *fru* and *dsx: sex-lethal (sxl), transformer (tra)*, and *transformer-2 (tra2)*. Whereas *dsx, sxl*, and *tra2* appear to be conserved at the sequence level in *D. virilis* (*2, 61, 62*), *tra* sequence comparisons suggest functional divergence in *D. virilis* and other *Drosophila* species (*63, 64*). Our findings here of divergent effects of FruM expression on sex-specific behaviors in *D. virilis* highlight the importance of going beyond sequence comparisons in carefully selected species for evaluating conservation vs. divergence of switch gene function.

In summary, through gene editing and careful behavior quantification, we found evidence for evolutionary divergence in *fru* function across *Drosophila* species. Future work should investigate species differences in *fru* circuitry underlying sex-specific behaviors.

## Materials and Methods

### Fly strains

We used *D. virilis* 15010-1051.47 (*41*) and *D. melanogaster* NM91 as wild-type strains. *D. virilis* were kept on standard medium at 20C on a 16:8 hr light:dark cycle (*65*) and aged 10-20 days, as this is the time required to reach sexual maturity (*66*). *D. melanogaster* were kept on standard medium at 25C on a 12:12 hr light:dark cycle and aged 3-7 days. Flies expected to produce male courtship behaviors (ie, males and *fru*^*∆ tra*^/*fru*^*∆ tra*^, *fru*^*∆ tra*^/*+*, and wild-type sibling (*D. virilis*) or control (*D. melanogaster*) females) were singly housed within 8 hours of eclosion, whereas courtship targets (wild-type females that were not siblings of *fru*^*∆ tra*^ females) were housed in groups of 5-6 flies.

### Comparison of *fruitless* nucleotide sequences

We downloaded the following data in April and May 2023: *D. melanogaster fru* (FBgn0004652) exon sequences from ensembl.org; the *D. virilis* scaffold (scaffold_12855) containing *fru* and the *D. simulans* chromosome (ch3R) containing most *fru* exons from the UC Santa Cruz Genome Browser (genome.uscs.edu); and the sequences containing the *D. simulans* B (accession number: GI: <u>111258132</u>) and C (accession number: <u>KF005597</u>) exons from Genbank (*67*). We used Geneious Prime 2023.1.2 to align the nucleotide sequence of each *D. melanogaster* exon to the relevant *D. virilis* and *D. simulans* sequences and recorded the % identity.

### Generation of *D. virilis fruitless* mutants *fru*^*∆ tra*^

To examine the role of *fruitless* in *D. virilis* song production, we used CRISPR-Cas9 mutagenesis to remove the Transformer (Tra) binding sites from the S exon, similar to the *fru*^*∆ tra*^ mutation previously made in *D. melanogaster* (*3*). We identified the Tra binding sites in *D. virilis* based on sequence similarity with *D. melanogaster* (*68*). We then designed CRISPR guide RNAs (gRNAs) flanking the Tra binding sites using the CRISPR Optimal target finder (*69*). The gRNAs had a 20-nucleotide target sequence and were flanked by a 3′ PAM sequence (‘NGG’) and a 5′ T7 RNA polymerase recognition sequence (‘GG’). The target genomic region was sequenced using Sanger sequencing. gRNAs are listed below. The 5’ is the T7 promoter, bold indicates the gRNA target, and italics indicate the 3’ portion that overlaps with the reverse primer. The protospacer adjacent mofit (PAM) is shown in parentheses.

L1: 5’ GAAATTAATACGACTCACTATA**GGTGTCTATGCCTAGGACTT**(AGG)*GTTTTAGAGCT AGAAATAGC 3’*

R1: 5’ GAAATTAATACGACTCACTATA**GGCTAGAGGCACGTGAGTAG**(TGG) *GTTTTAGAGCTAGAAATAGC 3’*

R2: 5’ GAAATTAATACGACTCACTATA**GGAACTGCATACCGTGCGGCA**(TGG) *GTTTTAGAGCTAGAAATAGC 3’*

The forward primer format and sgRNA-R primer sequences are based on (*70*). To generate the template for each sgRNA, we used the CRISPR forward and reverse 4 nmol Ultramer oligonucleotides (IDT). The full-length dsDNA template was amplified using Invitrogen Platinum PCR super mix high fidelity (Cat# 12532-016) and 0.5 µM forward and reverse primers. Reactions were carried on an Applied Biosystems 2720 Thermal Cycler, 95 °C 2 m, 35 cycles of (95 °C 20 s, 60 °C 10 s, 70 °C 10 s) and then purified with Ampure XP beads (A63880). In vitro transcription of 300 ng of sgRNA template DNA using T7 MEGAscript kit (Invitrogen AM1333) was carried out at 37 °C for 16-20h. Turbo DNase (Invitrogen AM2239) was added for an additional 15 min at 37 °C, then purified with Mega Clear Kit (Invitrogen AM1908). The gRNA concentration and quality were checked with Agilent Bioanalyzer and frozen in small aliquots at −80 °C for long-term storage. The CRISPR injection mixture contained 300 ng/µl recombinant Cas9 protein (PNA Bio CP01) and 40 ng/ µl sgRNA (per guide; we used one upstream L1 and two downstream R1 and R2 gRNA) and was injected into embryos of *D. virilis* wild-type strain 15010-1051.47. The Insect Transformation Facility at the University of Maryland performed all injections. We backcrossed the injected G0 flies to wild-type flies and selected lines in which germline mutagenesis was successful as determined by PCR genotyping (see below). PCR and sequencing confirmed that L1 and R2 successfully cut the DNA and removed 622bps, including the Tra binding sites. We obtained 12 independent lines carrying the *fru*^*∆ tra*^ mutant allele and observed no differences in behaviors across lines (data not shown).

### fruM-null

The method to remove the S-exon of the *fruitless* gene followed the same general procedure described for the *fru*^*∆ tra*^ design, with the following changes. We designed 2 CRISPR target sites (L-1 and L-3) upstream of the *D. virilis fru* S-exon start codon. The target sites downstream of the S-exon were as described earlier (R1 and R2).

L-1: GAAATTAATACGACTCACTATA**GGAAACCTTTAAACGGAGAAT** (TGG)*GTTTTAGAGCTAGAAATAGC*

L-3: GAAATTAATACGACTCACTATA**GGACCAACTAGTGCTAGAT**(CGG) *GTTTTAGAGCTAGAAATAGC*.

The injection mix contained the L-1, L-3, R1, and R2 gRNA. We crossed injected flies to one another and used PCR genotyping of the offspring to identify lines with germline transmission. PCR and sequencing confirmed that the L-1 and R1 guides successfully removed the S-exon. We obtained 7 independent lines in which the S-exon was removed.

### PCR genotyping

We used PCR to identify the genotype of experimental flies. We extracted DNA from the whole fly or from just the body (saving the heads for immunostaining from a subset of flies) using Quick-gDNA miniprep kit (Zymo Research R1050). We designed primers upstream and downstream of the Tra binding sites as follows:

CRISPRcut F3: TACGTACACGAATAGCCTCTTG

CRISPRcut R1: TGCCCGATTGAGCAAAATGC

We designed an additional primer upstream of the S-exon to identify flies in which the S-exon was successfully removed.

CRISPRcut F1: TGAGAGTTGTGTGATGGCTTG

### RT-PCR

We made total RNA from fly heads using Quick RNA Microprep kit (Zymo Research R1050). The reverse transcription reaction used the High Capacity cDNA Reverse Transcription Kit (Applied Biosystems 4368814) and RNase inhibitor (Promega N2515). We designed male-specific and female-specific forward RT-PCR primers, and a primer from a downstream *fru* common exon based on sequence similarity with those used previously in *D. melanogaster* (*8*).

Female-specific primer 10136 Fwd: GCAAAAGGAAGAGAGCCTCA

Male-specific primer 8200 Fwd: GATGGCCACCTCACAAGATT

Common primer C4 Rev: GCAGTCCATATTTCGAGACGA

### Immunohistochemistry

We dissected brains in ice-cold PBS, then fixed in 4% paraformadehyde in PBSX1 (Cellgro 21-040) with 0.3% Triton X-100 (Sigma Aldrich X100)(abbreviated PBT3) for 45 min in the dark at room temperature (RT). We blocked with 5% normal goat serum (NGS) (Life Technologies 16210-064) in PBT3 for two overnights at 4C. We incubated brains with 1:5000 rabbit anti-fruM (*33*) (gift from Stephen Goodwin) and 1:20 mouse anti-nc82 (Developmental Studies Hybridoma Bank, Iowa City, Iowa) in blocking solution for 3 overnights at 4C. We rinsed 8 times 20 min in PBT3 at RT before an overnight incubation in 1:500 goat anti-rabbit AlexaFluor 633 (Invitrogen A21070) and 1:500 goat anti-mouse AlexaFluor 488 (Life Technologies A11001) in blocking solution at 4C. We rinsed 4 times 20 min in PBT3, then 4 times 20 min in PBS at RT before leaving brains overnight in Vectashield (Vector Laboratories H-1000-10) at 4C. We mounted brains in Vectashield and imaged on a confocal microscope (Leica TCS SP8 X). Images were adjusted for brightness and contrast in Image J (NIH).

### Behavioral assays

Virgin males and females were used for behavior experiments. Video was captured at 60 frames/sec with a Point Grey Flea 3 CMOS camera (FL3-U3-13E4C-C). Audio was recorded at 10kHz on a 32-channel apparatus (*41, 71*). Each experimental fly was paired with a single homosexual +/+ partner on day 1 and with a single heterosexual +/+ partner on day 2. Each recording lasted 25 or 30 min. Experimental flies were singly housed to maintain their identity for PCR genotyping and/or checking for larvae. All assays occurred at ∼22C and between 0-4 hr ZT (*D. virilis*) or 0-1.5 hr ZT (*D. melanogaster*). Circular behavioral chambers were scaled for differences in fly body sizes between species: 11 mm diameter for *D. melanogaster* (*71*) and 20 mm diameter for *D. virilis* (*41*).

### Behavior quantification

We uniformly sampled each video and manually scored 10 sec bins 1 min apart for the presence or absence of the following behaviors: tarsal contact (front tarsi contact with any part of the other fly); proboscis extension; unilateral wing extension; bilateral wing extension; wing-flicking (unilateral wing extension to ∼45° that is quickly retracted without producing sound); low-posture fencing (flies face one another and extend their front tarsi toward each other while in a normal standing posture) (*43*); curling (female curls abdomen laterally with the tip of her abdomen pointed toward the male) (*44*); and spinning (female and male face in opposite directions and jointly move in a circle). Curling and spinning co-occurred so frequently that we did not attempt to score them separately. We report the percentage of bins containing at least one instance of the respective behavior in Fig 2B-D, Fig 4I-J, Supp Fig 3C-D, Supp Fig 4E-G, Q-R, and Supp Fig 5. We performed principal components analysis (PCA) on the amount of these 7 behaviors each fly produced when paired with a +/+ male and a +/+ female using Matlab 2019b. Copulation was defined as the time when the male first mounted the female and remained in the copulation position for at least 1 min in *D. virilis* (copulation duration ∼2-5 min) or 5 min in *D. melanogaster* (copulation duration ∼10-20 min) (*72*).

### *D. virilis* song segmenter

#### Network structure

The *D. virilis* song segmenter consisted of two convolutional neural networks in Python 3.4 (Supp Fig 2A-B): one network for classifying each point in the microphone recording as unilateral song, bilateral song, overlap of the two song types, or no song; and a second network for classifying bilateral song vs. no song. Using the 4-class network alone led to many bilateral song false positives, as noises such as grooming, jumping, rolling, etc, were often classified as bilateral song. Therefore, to improve the precision of bilateral song detection, we used a second, 2-class network to classify bilateral vs. no song. Both networks take the raw microphone recording as input. The four-class network used a window size of 4001 points (400.1 ms), batch size of 128, 6 epochs, 16 convolutional filters, 2x2 pooling, a convolutional kernel size of 9 convolutions and 4 padding, and trained on 10% of the data. The two-class network used a similar structure as the four-class network except it used a window size of 2001 points (200.1 ms) and 3 epochs during training.

#### Training data

A single observer used both audio and video to manually segment 5 30-min recordings of +/+ males paired with +/+ females, and 1 30-min recording of a +/+ male paired with a *fru*^*∆ tra*^/+ female. The *fru*^*∆ tra*^/+ female recording was chosen because of the large amount of song from both flies. We then drew from these data to make training data for the 2 neural networks. Training data for the 4-class network consisted of 31.6 total min (28 min from the *fru*^*∆ tra*^/+ and +/+ male pairing and the remaining from the +/+ male and female pairs), resulting in a total of 4.9% unilateral song (585 unilateral bouts), 25.4% bilateral song (10358 *fru*^*∆ tra*^/+ bilateral pulses, 92 +/+ bilateral pulses), and 1.3% overlap between unilateral and bilateral. We found the best performance when data from different rounds of data collection were included in the training set (data not shown). To make the training data for the two-class network, we removed unilateral and overlap portions from the training data for the four-class network, and added 21.5 min of additional noise and quiet taken from recordings of +/+ female pairs.

#### Song segmentation

To combine the output of the two segmenters, we first applied a median filter (window size of 10 ms) to the output probabilities and then averaged the bilateral and no song probabilities from the 4- and 2-class networks. We ignored the output of the 2-class network during portions identified as unilateral or no song by the 4-class network (Supp Fig 2C). The maximum probability at each timepoint was used to identify instances of song, with the following heuristics. To classify a point as bilateral song, we required the bilateral probability to be at least 1.25 times that of unilateral song, and to classify a point as overlap, we required the overlap probability to be at least 0.85. We required bilateral song within 40 ms before or up to 10 ms after unilateral song to have a classification probability of 1; otherwise it was assigned to unilateral song. We threw out segments predicted to be bilateral if they were shorter than 4 ms. We next defined unilateral bouts as contiguous unilateral predictions, and bilateral pulses as contiguous bilateral predictions. To separate contiguous unilateral or bilateral song into pulses, we first calculated the difference of the upper and lower peak envelopes of the microphone signal with a width of 5 ms, and then detected peaks in this signal. Minima on either side of these peaks were used to define where one pulse ended and the next started. We compared the segmenter output to ground truth data obtained by manually segmenting 22 recordings (Supp Fig 2D-F). Sensitivity, positive predictive value, and the harmonic mean (F) were calculated as previously described (*71*).

#### Song analysis

To calculate interpulse intervals (IPIs), we first calculated the difference of the upper and lower peak envelopes of the microphone signal with a width of 5 ms, and then detected peaks in this signal. The difference in timing of these peaks was taken as the IPI. Remaining song analysis was largely based on a previous study of *D. virilis* duets (*41*). A unilateral bout was defined as a series of at least 4 unilateral pulses with IPIs of 60 ms or less. A bilateral bout was defined as any number of bilateral pulses with IPIs of 100 ms or less. Unilateral response times were defined as the delay between the offset of a bilateral pulse and the onset of the immediately following unilateral bout. Bilateral response times were defined as the time between the onset of a unilateral bout and the center of the following bilateral pulse. Only response times less than 1.5 sec were included in our analysis.

For song feature measurements, we used only recordings with at least 10 unilateral and 10 bilateral pulses prior to copulation in copulating pairs, or over the entire recording otherwise. We quantified the amount of unilateral and bilateral song by summing the total time of the recording classified as unilateral or bilateral, then dividing by the courtship time, defined as the time between the first and last pulse (of either song type) in the recording (for non-copulating pairs) or the last pulse of either song type before copulation (for copulating pairs).

#### Statistics

Since most measurements were not normally distributed according to Jarque-Bera tests, we used non-parametric tests throughout. All statistics were performed in Matlab 2019b or 2022a.

## Acknowledgments

We thank Jan Clemens and Talmo Pereira for helpful discussions during the development of the *D. virilis* song segmenter; Stephen Goodwin and Michael Perry for anti-FruM antibody; Yun Ding, Michael Shahandeh, Zhilei Zhao, Duncan Mearns, and Max Aragon for comments on the manuscript; Darren Parker for discussions of *fruitless* exons; and members of the Murthy lab for feedback on analyses.

## Author contributions

Conceptualization: CAB, XJG, MM

Methodology: CAB, XJG, MC, MM

Investigation: CAB, XJG

Visualization: CAB

Supervision: MM

Writing - original draft: CAB, XJG, MM

Writing - review & editing: CAB, XJG, MC, MM

