## Supplementary Materials for "The role of *fruitless* in specifying courtship behaviors differs across *Drosophila* species"

Christa A. Baker *et al.*

**This PDF file includes:**

Figs. S1 to S5
Movies S1 to S5

**Other Supplementary Materials for this manuscript include the following:**
Movies S1 to S5

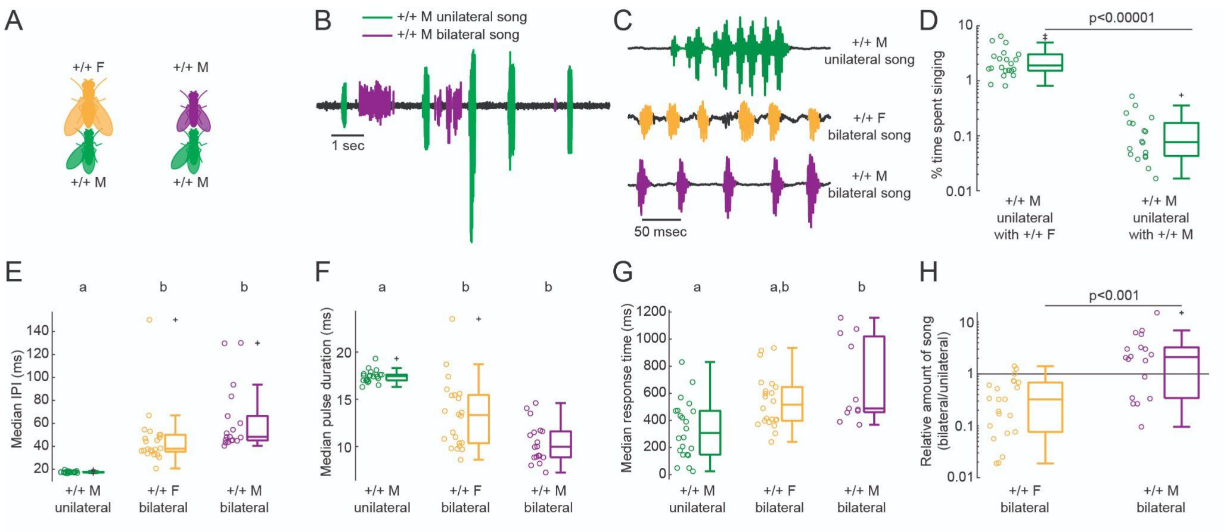


**Fig. S1.** ***D. virilis*** **bilateral song is sexually monomorphic.** (**A**) To compare bilateral song produced by +/+ males and females, we recorded song from heterosexual (left) and homosexual (right) pairs. (**B**) +/+ males are capable of singing bilateral song when courted by another male. (**C**) Bilateral song pulses in +/+ males are similar in appearance to +/+ female bilateral song and distinct from the stereotyped bouts of +/+ male unilateral song. (**D**) Total % of courtship time, defined as the time between the first and last pulse (from either sex) of each recording, that contains unilateral song. Although +/+ males take turns courting one another, the amount of male-directed courtship, as measured by the amount of unilateral song, is significantly lower than female-directed (Wilcoxon rank sum $z$=5.4, $p$<1e-7). (**E-G**) IPI (**E**), pulse duration (**F**), and response time (delay between onset of unilateral bout and center of first following bilateral pulse) (**G**) of +/+ male bilateral song compared to +/+ male unilateral and +/+ female bilateral songs. Each dot represents the within-fly median. Kruskal-Wallis and pairwise Wilcoxon rank sum tests were used to detect significant differences between groups (with Bonferroni correction). (**H**) +/+ males sing more total bilateral song than +/+ females relative to the amount of unilateral song in a given recording (Wilcoxon rank sum $z$=-3.7, $p$<1e-3). n=11-22 flies/group in (**D-H**).

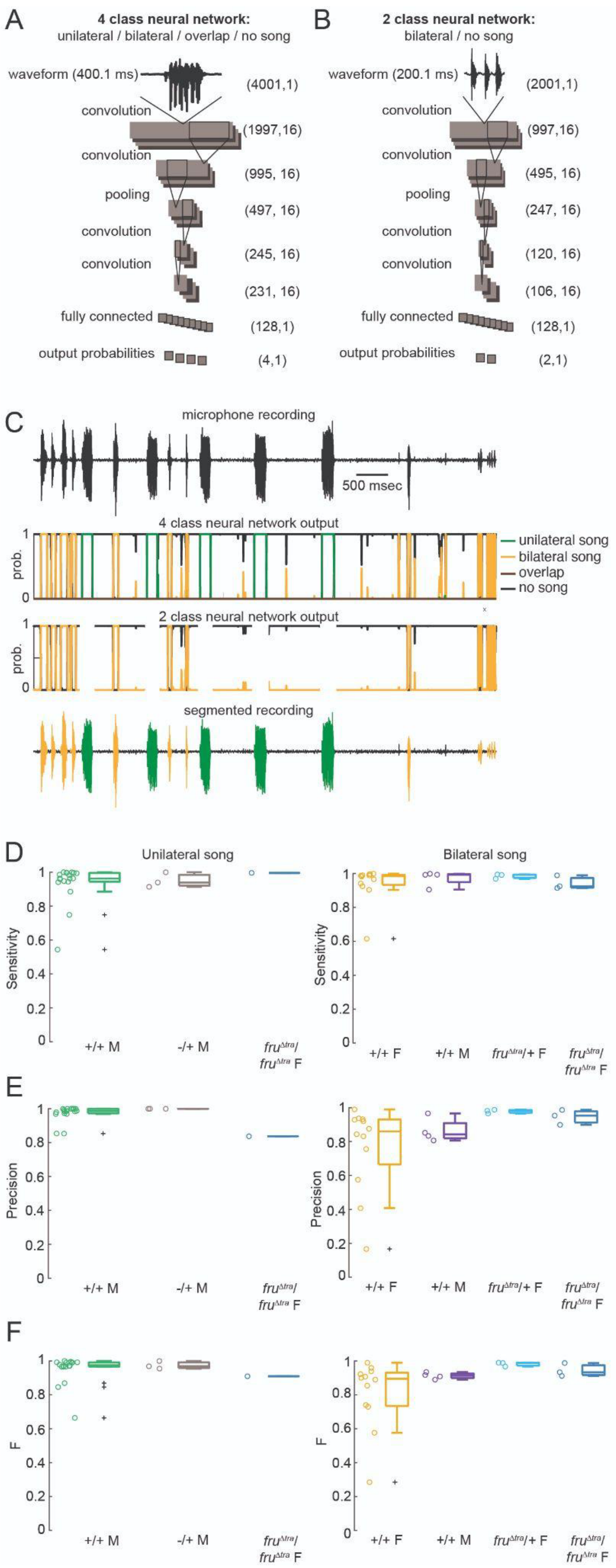
A
4 class neural network:
unilateral / bilateral / overlap / no song
waveform (400.1 ms)
(4001,1)
convolution
(1997,16)
convolution
(995, 16)
pooling
(497, 16)
convolution
(245, 16)
convolution
(231, 16)
fully connected
(128,1)
output probabilities
(4,1)
B
2 class neural network:
bilateral / no song
waveform (200.1 ms)
(2001,1)
(997,16)
(495, 16)
(247, 16)
(120, 16)
(106, 16)
(128,1)
(2,1)
C
microphone recording
500 msec
4 class neural network output
prob.
unilateral song
bilateral song
overlap
no song
2 class neural network output
segmented recording
D
Unilateral song
Bilateral song
Sensitivity
+/+ M
-/+ M
fru^Δtra/ fru^Δtra F
+/+ F
fru^Δtra/+ F
E
Precision
F
F

**Fig. S2. Convolutional neural network (CNN)-based song segmenter for *D. virilis* songs.** (**A**) Architecture of the CNN trained to distinguish between unilateral song, bilateral song, overlap of the two song types, and no song. The input is a raw microphone recording with a window size of 400.1 ms, and the output is a series of classification probabilities for each time point in the recording. This network was good at identifying unilateral song and overlap portions, but often classified noises, such as jumping or grooming, as bilateral song. (**B**) For this reason we trained a second network to specifically distinguish between bilateral song and no song. This 2-class CNN was similar to the 4-class network (**A**) except it used a smaller window size of 200.1 ms. (**C**) Example of song segmenter pipeline. Determinations of unilateral song and overlap (which is rare) come from the 4-class network, and thus those portions of the recording are ignored in the output of the 2-class network. The classification probabilities for bilateral song and no song are averaged between the two networks. The segmenter assigns each point in the recording according to the maximum classification probability, with a few heuristics (see Materials & Methods). (**D-F**) Sensitivity (**D**), precision (**E**), and their harmonic mean F (**F**) of the segmenter performance on unilateral (left) and bilateral (right) song compared to manual segmentations. n=18, 3, 1 (left) and 12, 4, 3, 3 (right) flies.

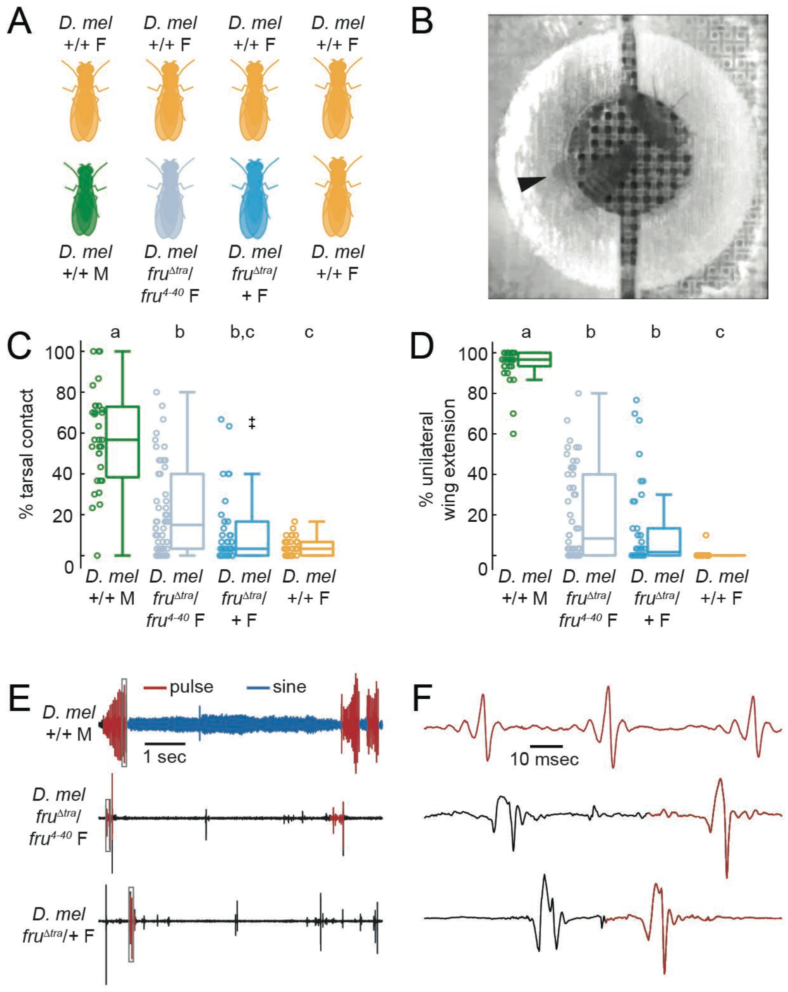


**Fig. S3.** ***D. melanogaster fru*$^{\Delta tra}$ females produce male courtship behaviors but aberrant unilateral song.** (**A**) We paired single *fru*$^{\Delta tra}$/*fru*$^{4\text{-}40}$ and *fru*$^{\Delta tra}$/+ *D. melanogaster* females with a +/+ (NM91) female. Single +/+ males and females each paired with a +/+ female served as controls. (**B**) A *fru*$^{\Delta tra}$/+ female performs unilateral wing extensions (arrowhead) directed toward a +/+ female. (**C-D**) Percentage of bins containing tarsal contact (**C**) or unilateral wing extensions (**D**) produced by each genotype when paired with a +/+ female. Each dot represents one fly. n=31, 54, 38, 34 flies. **(E)** Seven- second microphone recordings showing sounds concurrent with unilateral wing extensions directed toward a +/+ female. Unilateral wing extensions by males generate complex song bouts consisting of switches between pulse and sine song, whereas unilateral wing extensions by *fru*$^{\Delta tra}$ females only infrequently generate pulses (never sine). Some of these pulses are detected by the *D. melanogaster* song segmenter (red) (74). (**F**) Enlargements of the boxed regions in (**E**). *fru*$^{\Delta tra}$ female pulses detected by the *D. melanogaster* segmenter (red) lack the structure of +/+ male pulses (top).

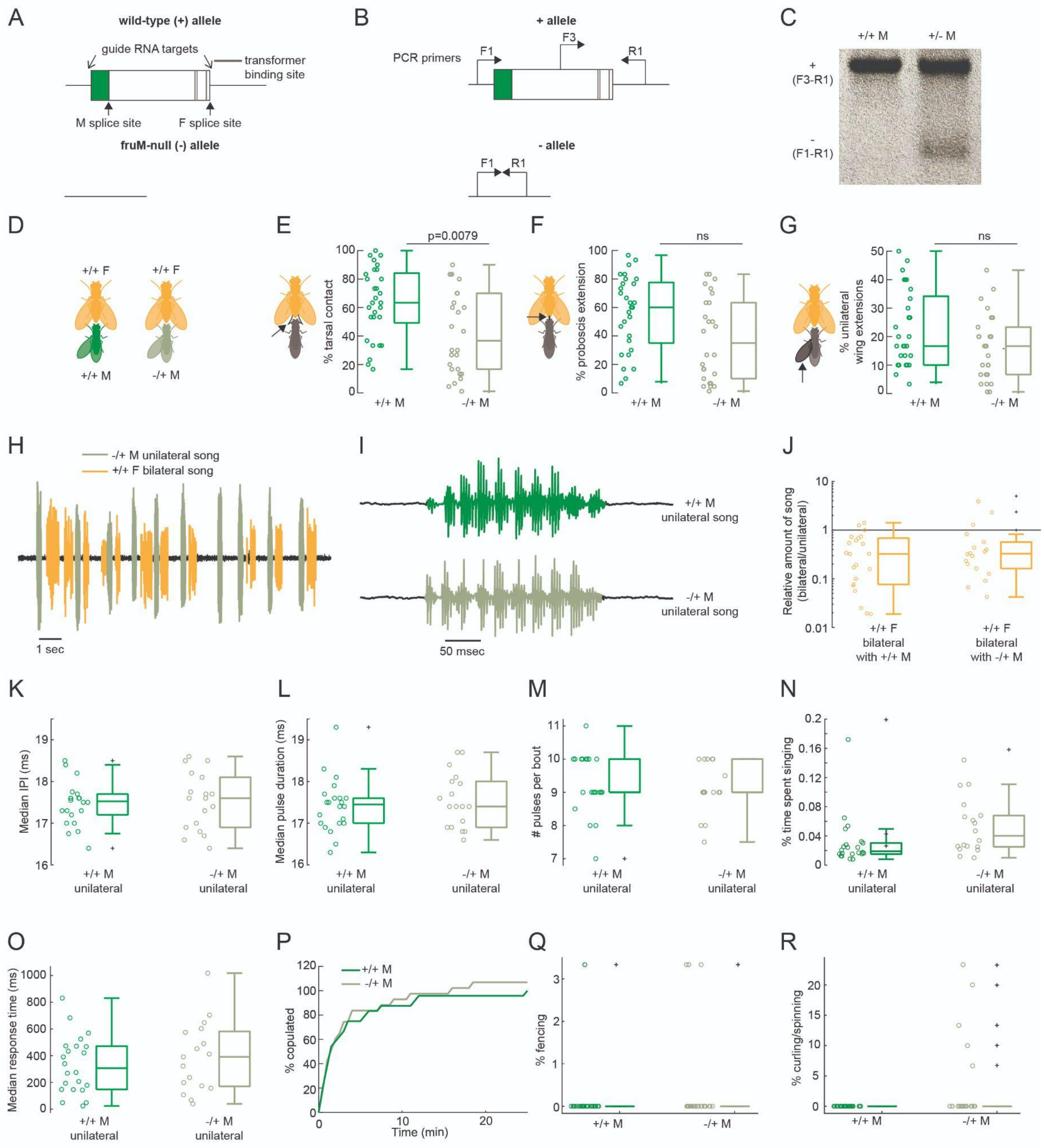


**Fig. S4. Removing a copy of *fruM* from *D. virilis* males has no effect on courtship behaviors.** (**A**) To generate a *fruM*-null allele, we designed CRISPR-Cas9 guide RNAs flanking the S-exon (top) to remove the S-exon (bottom). (**B**) We confirmed deletion of the S-exon with PCR using 1 reverse primer (R1) and 2 forward primers (F1, F3) due to the size of the S-exon. (**C**) PCR results using the primers shown in (**B**). In +/+ flies, the F1-R1 product is too large to amplify, so the product present is F3-R1 (left). The same product is present in -/+ flies due to the wild-type allele, in addition to the F1-R1 product, which is shorter due to the removal of the S-exon (right). (**D**) To test whether removal of a *fruM* copy affected male courtship behaviors, we paired single

-/+ males with a +/+ female. Single +/+ males (siblings to -/+ males) paired with a +/+ female served as controls. (**E-G**) Percent of bins containing tarsal contact (**E**), proboscis extension (**F**), and unilateral wing extension (**G**) directed toward a +/+ female. Each dot represents one fly. n=29 and 26 flies. (**H**) 14 sec microphone recording of a duet between a -/+ male and +/+ female. (**I**) A single unilateral song bout from a +/+ and a -/+ male. (**J-O**) Amount of bilateral song (from +/+ female) relative to unilateral song (from male) (**J**), median IPI (**K**), median pulse duration (**L**), median number of pulses per bout (**M**), percent time spent singing (**N**), and median response time (delay between offset of bilateral pulse and onset of following unilateral bout) (**O**) of unilateral song from each genotype when paired with a +/+ female. Each dot represents one fly. n=22 and 18 flies in (**J-O**). (**P**) Cumulative percent copulation over the 25 min observation period. Values are normalized relative to +/+ males. (**Q-R**) Percent bins with fencing (**Q**) and curling/spinning (**R**) behaviors. n=29 and 26 flies in (**P-R**).

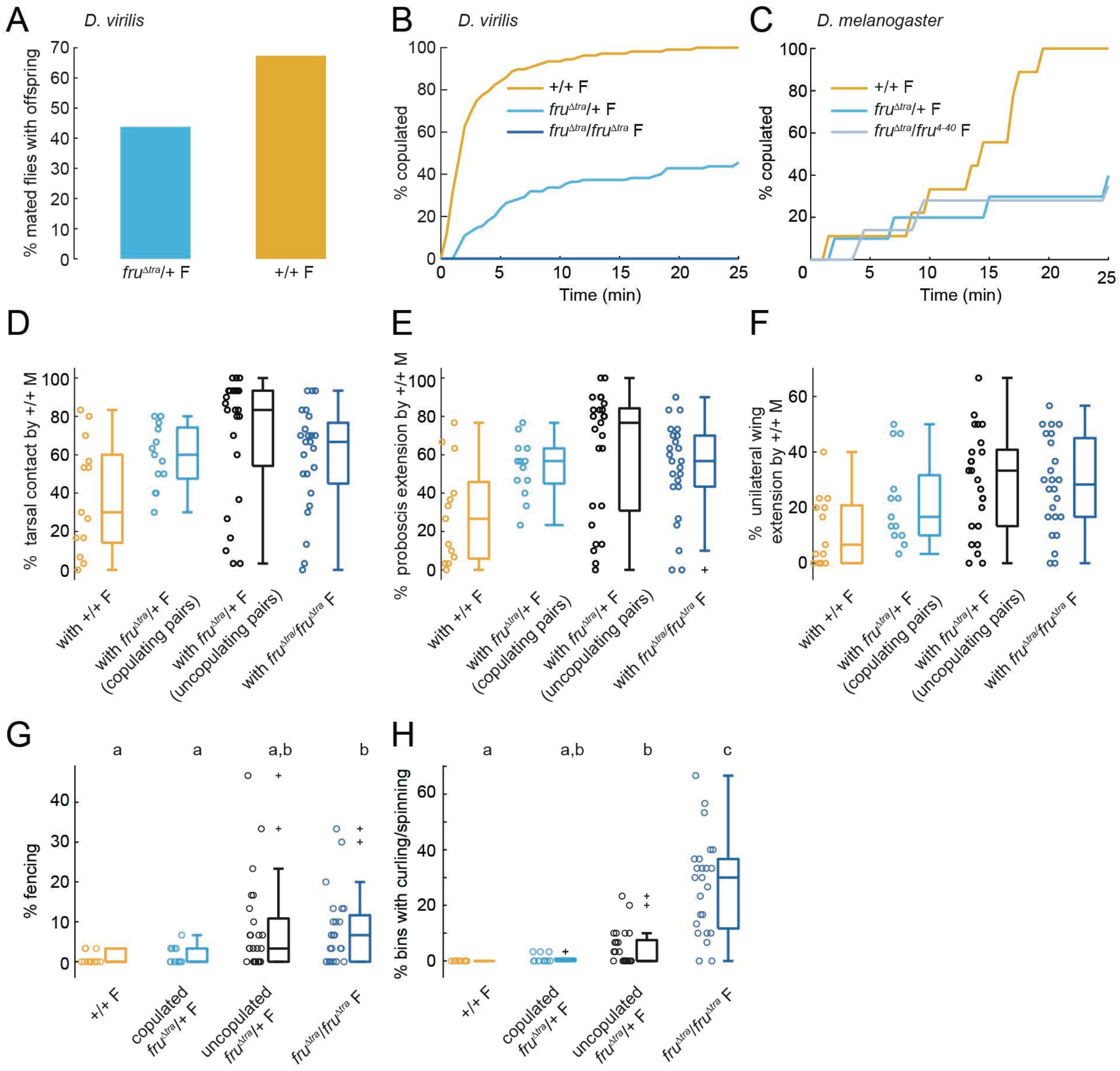


**Fig. S5. Effects of FruM expression on female reproductive behaviors of *D. virilis* and *D. melanogaster*.** (**A**) Percentage of matings with +/+ males that resulted in larvae in *D. virilis*. n=48, 107 flies. (**B**) Cumulative percentage of copulated *D. virilis* females (paired with +/+ conspecific males) over the 25 min observation period. Curves are normalized to +/+ females. n=118, 121, 24 flies. (**C**) Same as (**B**) for *D. melanogaster*. n=34, 38, 54 flies. (**D-F**) Percentage of bins containing tarsal contact (**D**), proboscis extension (**E**), and unilateral wing extension (**F**) from the +/+ male toward females of each genotype. Even within $fru^{\Delta tra}$/+ females, males produced equal amounts of courtship regardless of whether the pair copulated. Kruskal-Wallis one-way ANOVA tests were significant for all 3 variables (**D-F**) but pairwise Wilcoxon rank sum tests did not meet significance after Bonferroni correction for multiple comparisons. n=13, 13, 25, 24 flies. (**G-H**) Percentage of bins containing low-posture fencing (**G**) and curling/spinning (**H**) between females paired with +/+ males. There were no differences in the amount of these two behaviors in copulating vs. non-copulating *fru*$^{\Delta tra}$/+ females. n=13, 13, 25, 24 flies. Statistical tests were Kruskal-Wallis one-way ANOVA followed by pairwise Wilcoxon rank sum tests with Bonferroni correction.

**Movie S1.** *D. virilis* $fru^{\Delta tra}/fru^{\Delta tra}$ female taps, licks, and sings unilateral song to a +/+ female (white dot). The +/+ female responds with bilateral song.

**Movie S2.** *D. melanogaster* $fru^{\Delta tra}/+$ female taps, licks, and performs unilateral wing extensions directed toward a +/+ female (white dot). The wing extensions produce some sounds but not male-typical song.

**Movie S3.** *D. virilis* $fru^{\Delta tra}/fru^{\Delta tra}$ female sings bilateral song while duetting with a +/+ male (smaller fly).

**Movie S4.** Low-posture fencing between *D. virilis* $fru^{\Delta tra}/fru^{\Delta tra}$ female (larger fly) and +/+ male. The interaction begins with duetting, then the $fru^{\Delta tra}/fru^{\Delta tra}$ female turns to the male and approaches him with her front tarsi. She also produces a brief curling event and a few wing flicks. The male quickly resumes courting her once the aggression subsides.

**Movie S5.** Curling and spinning between *D. virilis* $fru^{\Delta tra}/fru^{\Delta tra}$ female (larger fly) and +/+ male. The interaction begins with the male courting the $fru^{\Delta tra}/fru^{\Delta tra}$ female and then the $fru^{\Delta tra}/fru^{\Delta tra}$ female initiates curling, in which she curls her abdomen toward the male and at times shoves him with it. The male appears to try to continue tapping and licking the $fru^{\Delta tra}/fru^{\Delta tra}$ female from behind, which results in the pair spinning around together. Duetting immediately resumes when the spinning bout ends.
